# Ecological drift during colonization drives within- and between-host heterogeneity in animal symbiont populations

**DOI:** 10.1101/2023.08.21.554070

**Authors:** Jason Z. Chen, Zeeyong Kwong, Nicole M. Gerardo, Nic M. Vega

**Affiliations:** Department of Biology, Emory University, Atlanta, GA USA; Laboratory of Bacteriology, National Institutes of Allergy and Infectious Diseases, Hamilton, MT USA; Department of Physics, Emory University, Atlanta, GA USA

## Abstract

Host-associated microbiomes vary greatly in composition both within and between host individuals, providing the raw material for natural selection to act on host-microbe associations. Nonetheless, the drivers of compositional heterogeneity in host-associated microbiomes have only rarely been examined. To understand how this heterogeneity arises, we utilize the squash bug, *Anasa tristis*, and its bacterial symbionts in the genus *Caballeronia*. We artificially modulate symbiont bottleneck size and strain diversity during colonization to demonstrate the significance of ecological drift, which causes stochastic fluctuations in community composition. Consistent with predictions from the neutral theory of biodiversity, ecological drift alone can account for heterogeneity in symbiont community composition between hosts, even when two strains are nearly genetically identical. When acting on competing, unrelated strains, ecological drift can maintain symbiont genetic diversity among different hosts by stochastically determining the dominant strain within each host. Finally, ecological drift mediates heterogeneity in isogenic symbiont populations even within a single host, along a consistent gradient running the anterior-posterior axis of the symbiotic organ. Our results demonstrate that symbiont population structure across scales does not necessarily require host-mediated selection, but emerges as a result of ecological drift acting on both isogenic and unrelated competitors. Our findings illuminate the processes that might affect symbiont transmission, coinfection, and population structure in nature, which can drive the evolution of host-microbe symbiosis and microbe-microbe interactions within host-associated microbiomes.

## Introduction

All multicellular life forms on Earth play host to teeming communities of host-associated microbes. The most specialized host-microbe associations, or symbioses, have evolved multiple times in the tree of life. Host-microbe symbioses feature diverse mechanisms for host benefit, control of community composition, and limited opportunities for microbial transmission [1]. These mechanisms play essential roles in initiating, stabilizing, and undermining symbiosis during a host’s lifetime, over generations, and across macroevolutionary timescales [1,2]. Nevertheless, a persistent paradox in the study of host-microbe symbioses is that, like microbes in natural environments, microbial symbionts exhibit enormous strain diversity [3–10], even when natural selection, imposed by specialized interactions with their hosts, is expected to erode genetic variation.

Different mechanisms, based on environmental or host-level selection, are typically invoked to explain the maintenance of symbiont genetic variation, often in terms of host benefit [11,12]. However, these hypotheses do not account for how host-associated consortia assemble as ecological communities, which embeds this genetic variation within patches in physical space [13,14]. This is an inherently stochastic process that generates heterogeneity [15–17]. Heterogeneity in host-associated microbial communities manifests at two scales: as heterogeneity in colonization *between hosts*, and as spatial heterogeneity across tissues and organs *within each host*. At both scales, it is critical to understand how heterogeneity emerges during establishment of symbiosis, which drives the evolution, ecology, and physiology of both host and microbe.

While the ecological processes that create heterogeneity during community assembly have been studied with mathematical models (e.g. [18]), validation of these models in empirical studies using natural, ecologically realistic communities, including host-associated microbial communities [14–16,19], is scarce. Some of these processes are deterministic, acting on specific traits that allow or hinder establishment of a taxon in a predictable, niche-based fashion. However, community assembly is also governed by dispersal between habitats. Dispersal imparts a stochastic element on community assembly [16]: Taxa immigrate and establish in new patches in a probabilistic manner, in part because they experience transient reductions, called bottlenecks, in population size [20]. These bottlenecks intensify ecological drift, or stochastic variation in community composition. Since the proposal of Hubbell’s unified neutral theory of biodiversity, the relative role of stochastic processes such as ecological drift in community assembly, compared with deterministic niche-based processes such as between-species interactions, has been a matter of continuous study [21–23].

To illustrate the importance of ecological drift during the establishment of even highly specific symbioses, we employ the squash bug, *Anasa tristis* (Fig 1A), as a model. *A. tristis* is host to specific symbionts in the β-proteobacterial genus *Caballeronia* [24] (previously referred to in the literature as the *Burkholderia* “SBE” clade [25] or the *B. glathei*-like clade [26]), which it requires for survival and normal development to adulthood. Acquisition of *Caballeronia* occurs through the environment after nymphs (immature insects) molt into the second instar. Once they successfully colonize the host, symbionts are housed in hundreds of sacs called crypts, which form two rows running along a specific section of the midgut, called the M4 (Fig 1B). Unlike many other insect symbionts, *Caballeronia* can be isolated from bugs and established in pure culture in the laboratory. Because *A. tristis* nymphs hatch from the egg symbiont-free, the symbiosis can be reconstituted anew every generation by feeding cultured symbionts to these nymphs in the laboratory.

**Fig 1.**
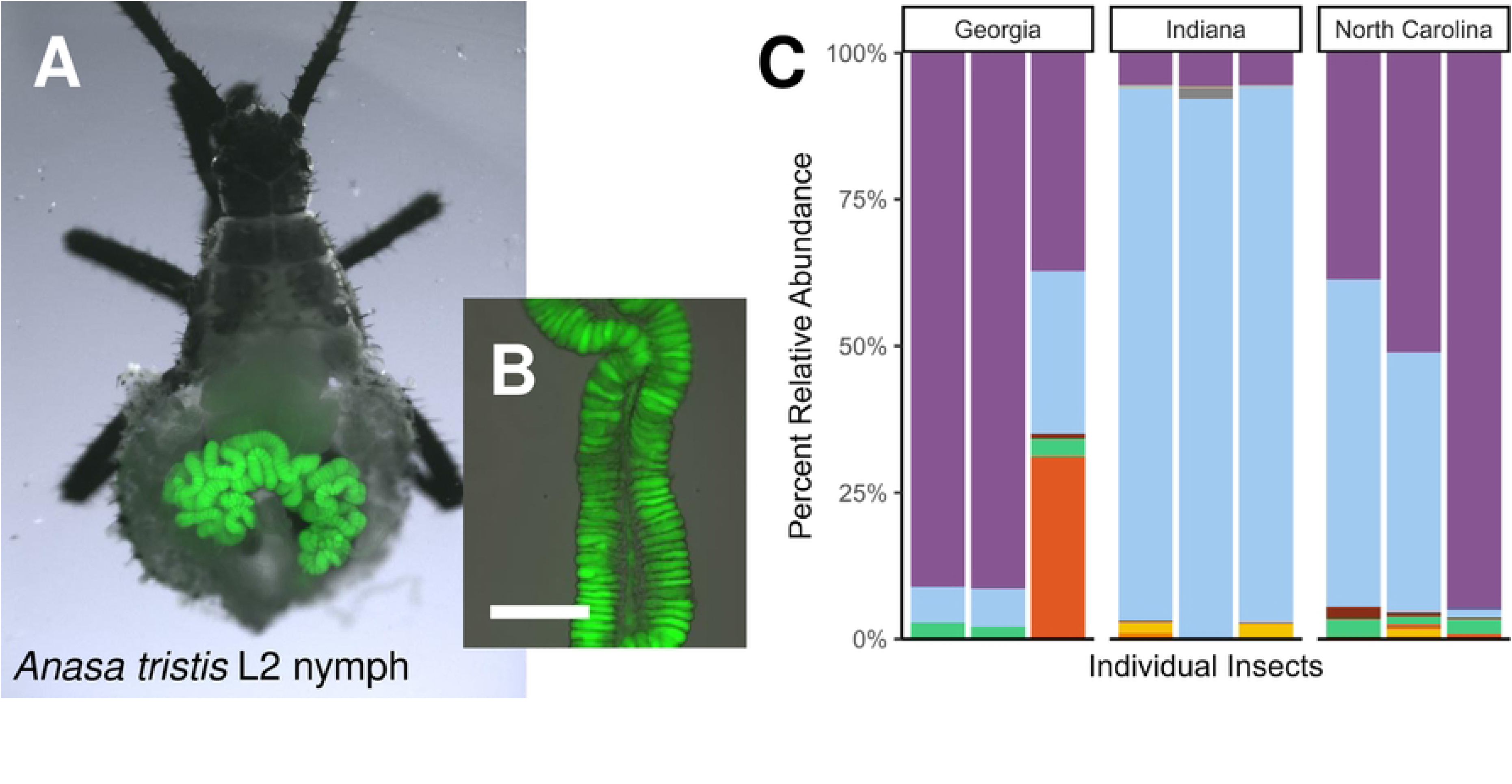
The squash bug, *Anasa tristis*, engages in a specialized host-microbe symbiosis with bacteria in the genus *Caballeronia*. A) A second instar squash bug (*Anasa tristis*) dissected to reveal the M4 section of the midgut, colonized with *Caballeronia* symbionts expressing the sfGFP fluorescent protein. B) Inset: The fine structure of the M4, consisting of two rows of hundreds of sac-like crypts lining a central lumen. Crypts are colonized at high density with *Caballeronia* symbionts expressing the sfGFP fluorescent protein. The scale bar represents 250 µm. C) Relative abundance of the top 20 *Caballeronia* 16s V3-V4 amplicon sequence variants (ASVs) within 9 bugs across 3 field localities in Georgia, Indiana, and North Carolina, with different colors representing different ASVs. Modified from Stoy *et al.* (2023) [9].

Because the host relies on *Caballeronia* strains as its symbiotic partners, its digestive tract imposes strong selection to favor *Caballeronia* colonization [27,28], similar to other specialized systems [29]. As a result, *Caballeronia* constitutes the vast majority of the microbial community within the M4 symbiotic organ, even though squash bug nymphs are exposed to diverse environmental microbes on squash fruit and plants [24]. However, this and similar bug-*Caballeronia* symbioses are extremely non-specific below the genus level [30], with different bug isolates conferring nearly the same degree of host benefit across different *Caballeronia* clades [9,31]. In accordance with this apparent lack of specificity, we observe that within-host communities from wild insect populations vary widely in their composition [9] (Fig 1C). So, beyond the coarse ecological filter that the host insect applies against non-symbiotic taxa [27,28], little is known about the ecological processes that maintain within- and between-host diversity of this beneficial symbiont.

Here, we explore the hypothesis that both within- and between-host diversity in symbiont populations arise stochastically as a result of ecological drift during infection [16]. First, we set out to explore a range of conditions under which this pattern might emerge, incorporating neutral competition (where all cells are isogenic, and thus functionally equivalent, individuals) [22] and interspecies competition (where cells are genetically distinct, but still equally host beneficial) between symbiont strains. By experimentally manipulating transmission bottleneck size, we show that ecological drift alone can account for heterogeneity between hosts, segregating strains between hosts and decreasing the probability of coinfection. Using isogenic coinfections, we additionally demonstrate that the symbiotic organ imposes spatial heterogeneity on within-host populations, whereby separate crypts are colonized by different strains. Our results demonstrate the role of ecological drift in the assembly of a highly specialized host-microbe system, and point out its significance in structuring symbiont population diversity across scales.

## Results

### Ecological drift is sufficient to generate variation in colonization outcome

We reasoned that if ecological drift plays a role in generating heterogeneity in symbiont populations between hosts, it should generate greater and greater heterogeneity under smaller and smaller inoculum densities, which represent tightening transmission bottlenecks in our experiments. Specifically, according to the neutral model [22], under tight bottlenecks, colonization outcomes should be bimodal, with hosts dominated by single isolates. To test this hypothesis, we implemented a simple experimental design (Fig 2A), previously applied to human pathogens, that modulates transmission bottleneck size while maintaining the relative abundance of each strain during transmission [32]. To minimize the involvement of selection, we used isogenic, green- and red-fluorescently labelled isolates of *C. zhejiangensis* GA-OX1, a highly beneficial strain isolated previously from *A. tristis* [9]. Because our experiments involved only two competitors, we use the bimodality coefficient [16,33], among other measures, to quantify heterogeneity in community composition. We inoculated second instar squash bug nymphs with approximately 1:1 mixtures of GA-OX1 sfGFP with GA-OX1 RFP, diluted to produce inocula ranging from approximately 10^6^ to 10^1^ CFU/µL (Fig S1A).

**Fig 2.**
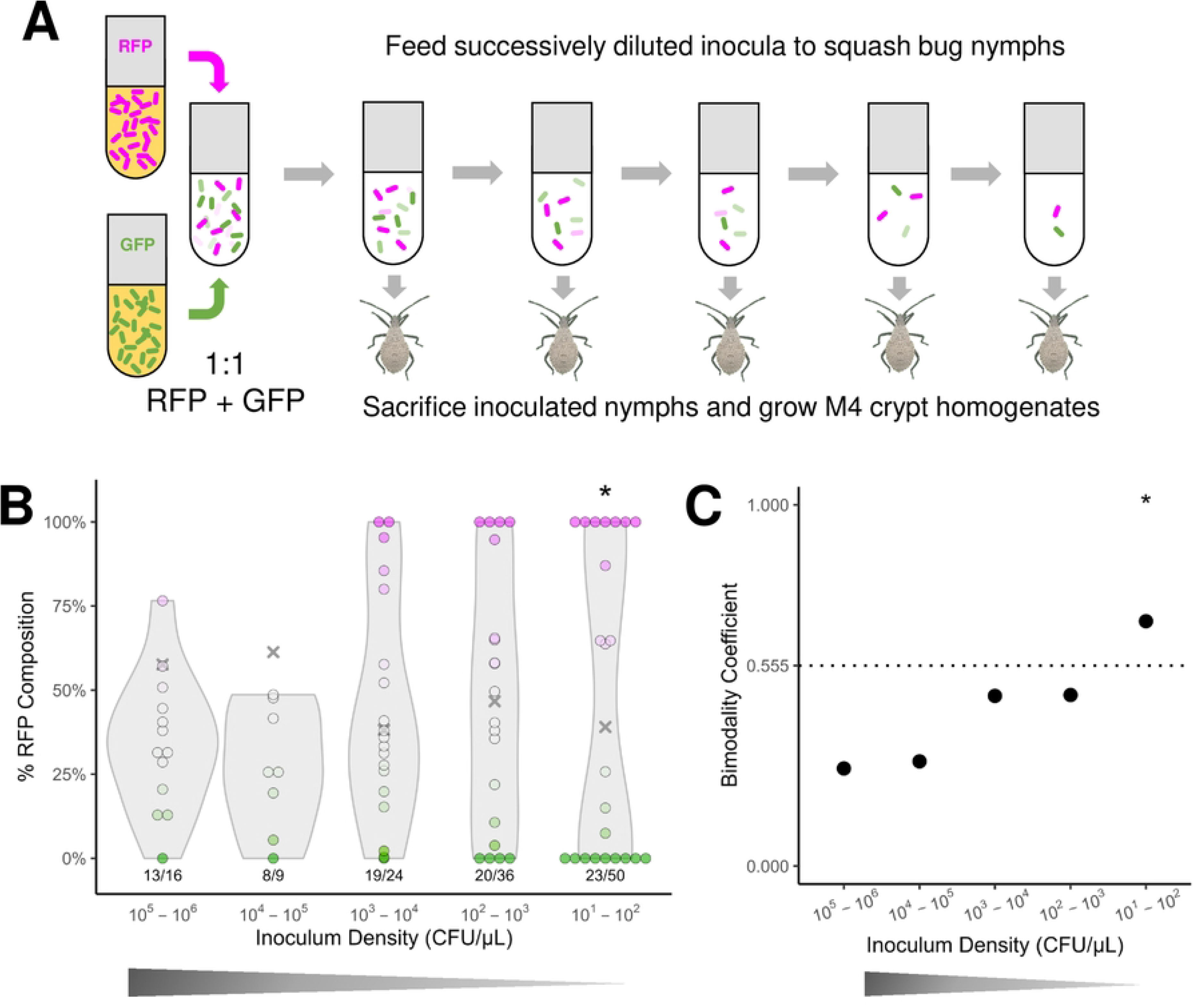
The strength of ecological drift mediates variability in the outcome of symbiont colonization. A) Experimental design. Symbionts previously isolated from squash bugs, made to stably express green or red fluorescent proteins (GFP or RFP), are grown individually in liquid culture. Liquid cultures are combined at a pre-determined ratio, then the mixture is diluted at various concentrations in the inoculation medium such that the inoculum density (a proxy for transmission bottleneck size) varies over several orders of magnitude while retaining the relative abundance of each strain across inocula. B) Variable colonization outcomes associated with different transmission bottleneck sizes in isogenic co-inoculation, using *Caballeronia zhejiangensis* GA-OX1 sfGFP and RFP. Gray X marks indicate the mean % GA-OX1 RFP associated with each inoculum treatment, ranging from 101 to 106 CFU/µL. Points represent individual nymphs, and the color of each point and its position along the y-axis represent the percent relative abundance of GA-OX1 RFP colonies among all fluorescent colonies recovered from each nymph. Magenta points represent nymphs from which only RFP colonies were recovered, green points represent nymphs from which only GFP colonies were recovered, and faded magenta/green colonies represent coinfected nymphs. Violin plots associated with each treatment depict the shape of the distribution in relative RFP abundance. Asterisks indicate significantly multimodal infection outcomes as determined by Hartigan’s dip test, at a significance value of p < 0.05. Below each violin plot, the success rate of colonization is indicated as the number of nymphs that were successfully colonized with *Caballeronia* out of all nymphs sampled. Trials were aggregated across multiple runs. C) Bimodality coefficients calculated from results in panel B. The 0.555 threshold (marked with a dotted line) indicates the bimodality coefficient expected from a uniform distribution. Asterisks indicate significantly multimodal infection outcomes as indicated by Hartigan’s dip test, at a significance level of p < 0.05.

Under the highest inoculum densities, corresponding to the loosest bottlenecks, differences between the M4 communities of individuals are minimized, with a slight bias in favor of the sfGFP strain (Figs 2B and S1B). The slight bias towards sfGFP colonization could be due to toxic aggregation of the dTomato fluorescent protein, which has been observed in eukaryotic cells [34]. As inoculum density decreases, and thus as transmission bottlenecks tighten, individual infections become increasingly dominated by one or the other strain, causing the bimodality coefficient to increase (Figs 2C and S1C, Table S1). Below 100 CFU/µL, individual infections are comprised of mostly either GFP or RFP, manifesting as a weakly but significantly bimodal outcome (bimodality coefficient = 0.677, Hartigan’s dip statistic = 0.152, p<2.2e-16) (Table S1). Fluorescence images of whole nymphs provided qualitative confirmation of our results, with individual insects exhibiting heterogeneity in RFP and GFP fluorescence at lower inoculum densities (Fig S2). Through this set of experiments, we show that ecological drift is sufficient to drive heterogeneous colonization outcomes.

### Ecological drift maintains coexistence between competing strains in separate hosts

Having illustrated the action of transmission bottlenecks on a single symbiont genetic background, we sought to understand next how they would act on genetically distinct host-beneficial strains. If ecological drift has an effect even when selection can act on competitive differences between strains, we should see a similar result to our previous experiment, with bimodality increasing with tightening transmission bottlenecks. We tested *C. zhejiangensis* GA-OX1 alongside *C. sp. nr. concitans* SQ4a [24], which represent two lineages within the *Caballeronia* genus (Fig S3) [26,35] but are nonetheless equally beneficial for host developmental time and survivorship in the laboratory[9,24]. SQ4a was previously labelled with GFPmut3 [24,36], and was additionally labelled with sfGFP and dTomato for this study using the same constructs [37] that were applied to GA-OX1 above.

First, we demonstrated that GA-OX1 and SQ4a compete under an *in vitro* approximation of natural conditions within the host midgut (Fig 3A). In trials where SQ4a sfGFP and RFP were grown together as liquid cultures in filter-sterilized zucchini squash extract, both strains were recovered at high densities after 24 hours. On the other hand, when either SQ4a strain was grown with a counter-labelled GA-OX1, SQ4a almost always went extinct (T-test, p<0.001, n=10). Labelled GA-OX1 strains grew to high densities regardless of whether they were growing alongside SQ4a or the counter-labelled GA-OX1. These data suggest that GA-OX1 is the superior competitor to SQ4a *in vitro*.

**Fig 3.**
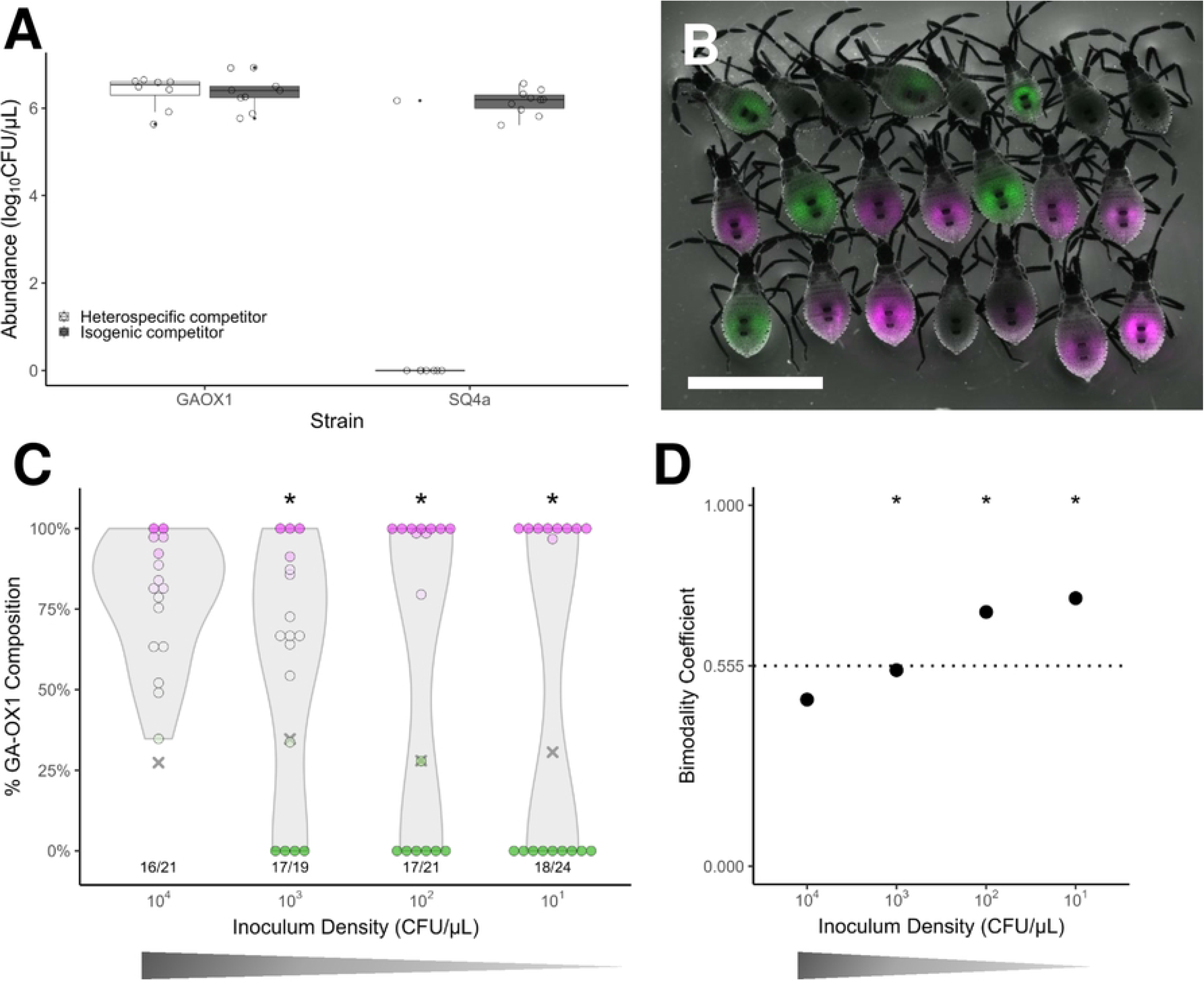
Ecological drift mediates heterogeneity in host infection by competing strains. A) Competitive interactions between two symbiont strains in liquid culture. GA-OX1 and SQ4a differentially labelled with sfGFP or RFP were cocultured with a strain with the opposite fluorescent marker, either of an isogenic background or of the other species, then assayed after 24 hours for abundance. White boxplots represent growth in the presence of a heterospecific competitor, while gray boxplots represent growth in the presence of an isogenic competitor. B) Fluorescence image of one cohort of squash bug nymphs fed a combination of SQ4a sfGFP and GA-OX1 RFP, at a combined density of ∼5000 CFU/µL. Green indicates the presence of SQ4a sfGFP in nymphs, and magenta indicates the presence of GA-OX1 RFP. Nymphs without fluorescence were not successfully colonized with either symbiont strain. Scale bar indicates 5 mm. C) Variable colonization outcomes associated with different transmission bottleneck sizes in two-species co-inoculation, using *C. sp. nr. concitans* SQ4a GFPmut3 and *C. zhejiangensis* GA-OX1 RFP. Gray X marks indicate the percent GA-OX1 RFP associated with each inoculum treatment, ranging from 10^1^ to 10^4^ CFU/µL. Points represent individual nymphs, and the color of each point and its position along the y-axis represent the percent relative abundance of GA-OX1 RFP colonies among all fluorescent colonies recovered from each nymph. Magenta points represent nymphs from which only GA-OX1 RFP colonies were recovered, green points represent nymphs from which only SQ4a GFPmut3 colonies were recovered, and faded magenta/green points represent coinfected nymphs. Violin plots associated with each treatment depict the shape of the distribution in relative GA-OX1 RFP abundance. Below each violin plot, the success rate of colonization is indicated, as the number of nymphs that were successfully colonized with *Caballeronia* out of all nymphs sampled. Asterisks indicate significantly multimodal infection outcomes as determined by Hartigan’s dip test, at a significance level of p < 0.05. D) Bimodality coefficients calculated from results in panel C. The 0.555 threshold (marked with a dotted line) indicates the bimodality coefficient associated with a uniform distribution. Asterisks indicate significantly multimodal infection outcomes as indicated by Hartigan’s dip test, at a significance value of p < 0.05.

We next co-colonized hosts with mixtures of GA-OX1 RFP and SQ4a GFPmut3, using the same experimental design as in single-strain colonization. As expected, high inoculum density (≥10^4^ CFUs) biased infections in favor of the superior competitor, GA-OX1 (Fig 3C). Decreasing inoculum densities resulted in strong bimodality in infection outcomes, where individual hosts were dominated either by the superior competitor GA-OX1 or by the weaker competitor SQ4a (Fig 3C, Table S2). This bimodality was qualitatively reproducible across different combinations of SQ4a and GA-OX1 expressing different fluorophores from different synthetic constructs (Figs 3B and S4, Table S3 and S4). Bimodality coefficients were consistently higher at high colonization densities in interspecific competition experiments than neutral competition experiments (Figs 3D, S4B, and S4D, Tables S2-S4).

### Ecological drift during colonization generates within-host spatial heterogeneity

The squash bug symbiotic organ, called the M4, contains hundreds of crypts (Figs 1A and 1B). Because each crypt is filled with its own population of symbionts, we asked whether these crypts might exhibit heterogeneity in symbiont composition within the host, consistent with previous unquantified observations from related insect-*Caballeronia* models [27,38]. If crypts indeed contain heterogeneous populations, we would expect crypts to contain mostly RFP- and mostly GFP-expressing symbionts, as opposed to highly similar populations composed of one or both types. We also examined if within-host heterogeneity might be sensitive to inoculum density in the same manner as between-host heterogeneity. If so, we would expect individual crypts to be more heterogeneous when symbionts are subjected to tighter transmission bottlenecks during host colonization.

We systematically characterized within-host spatial heterogeneity by co-inoculating nymphs with 1:1 mixtures of counter-labeled GA-OX1 at approximately 10^6^ and 10^2^ CFU/µL, as above. By imaging freshly dissected whole guts from coinfected nymphs, we observed that the M4 does impose spatial heterogeneity on symbiont populations, with individual crypts varying in GFP and RFP intensity even at colonization with 10^6^ symbiont CFU/µL (Fig 4A). However, there is a clear gradient in the degree of heterogeneity among crypts along the length of the M4, with anterior crypts being co-colonized and posterior crypts being singly infected (Fig 4B). We quantified this gradient by measuring the variance in RFP intensity relative to GFP along the length of the M4 (Fig 4C). Contrary to our expectations, we saw that nymphs colonized with just 10^2^ symbiont CFU/µL also exhibited this gradient, with anterior crypts being co-colonized despite a 10,000-fold reduction in inoculum density (Figs 4B and 4C). Thus, patterns of heterogeneity within the host are consistent over four orders of magnitude in inoculum density. Even when microbe-microbe competition is nearly neutral, host anatomy appears to impose spatial structure on symbiont populations.

**Fig 4.**
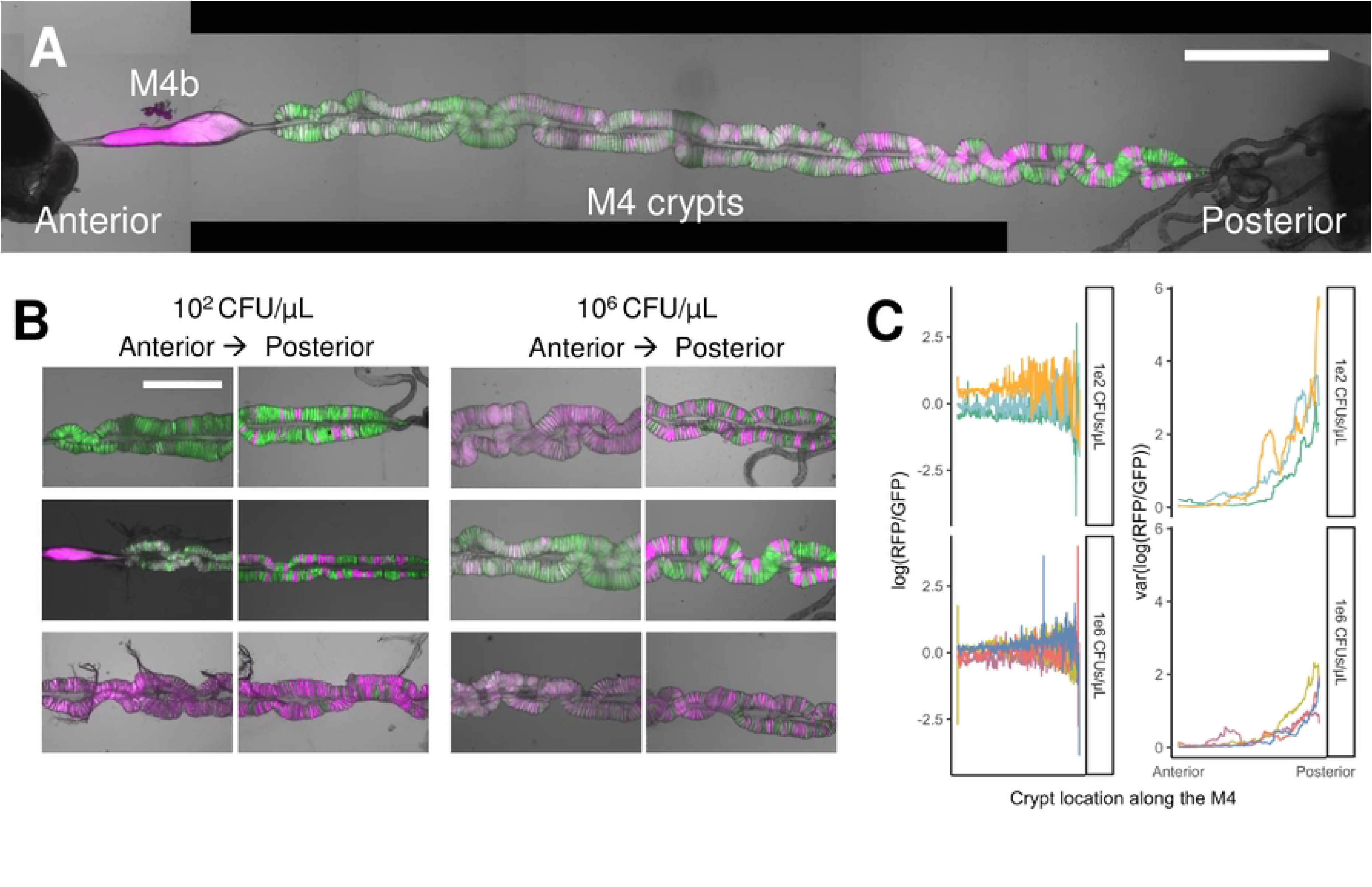
The squash bug symbiotic organ (the M4) imposes spatial heterogeneity on *Caballeronia* populations within the host. a) The entire M4 of a representative second instar nymph, inoculated with a combined 10^6^ CFU/µL GA-OX1 GFP and RFP, dissected and linearized to illustrate symbiont colonization along its length. The anterior end of the M4 is oriented to the left and the posterior end is oriented to the right. The magenta, spindle-shaped organ is the M4b, which is functionally distinct from the crypts that house the symbiont population. The scale bar represents 1 mm. b) Anterior and posterior crypts from 3 nymphs fed 10^2^ CFU/µL (left) and 3 other nymphs fed 106 CFU/µL (right) GA-OX1 GFP and RFP, dissected and prepared as in a). For each specimen, the anterior crypts are on the left and the posterior crypts are on the right. In each image, specimens are oriented as in a), with the anterior to the left and the posterior to the right. The scale bar represents 500 µm. c) Ratio of normalized RFP intensity relative to normalized GFP intensity (left) and variance in this ratio within a sliding window (right) along a transect from the anterior to the posterior of the M4. Nymphs were either inoculated with 10^2^ CFU/µL (n=3, top) or 10^6^ CFU/µL (n=4, bottom), and different colored lines represent the trajectories of these values associated with each nymph.

## Discussion

Previous research on a suite of closely related insect-*Caballeronia* symbioses has uncovered both heterogeneity in symbiont composition and low diversity of symbiont populations within hosts between hosts [9,30,39,40]. In the present study, we reveal the processes that underlie these patterns are consistent with stochastic colonization, which results in strong ecological drift as symbionts establish in their host insects. By modulating transmission bottleneck sizes of inocula containing isogenic, nearly neutrally competing strains, we show that ecological drift alone can generate heterogeneity in colonization outcome between different hosts [15,16], consistent with the neutral theory of biodiversity [22,23]. Next, by manipulating bottleneck sizes of inocula containing different symbiont species, we not only experimentally demonstrate the role of ecological drift in maintaining genetic diversity but also highlight the role of between-symbiont competition. Ecological drift generates variation in founding populations between hosts, while competition drives homogeneity in symbiont populations during subsequent proliferation inside a single host. The effect is bimodality in symbiont colonization, even when transmission bottlenecks are loose. Our results mirror findings from similar studies using plant communities, where competitive asymmetries between species also exaggerate the effect of ecological drift [21], and call attention to the role that intersymbiont competition might play during the early stages of colonization in other host-microbe systems [15,41–43].

Although we implemented our experiments using horizontal transmission, many symbioses exhibit elaborate, host-controlled mechanisms that ensure vertical transmission from parent to offspring [2,44–47], including even our own model system [48]. We argue that vertical transmission strategies are not immune to the effects of ecological drift, which acts on communities regardless of how they disperse. Indeed, some vertically transmitted symbionts undergo extreme transmission bottlenecks [49–51], exaggerating the intensity of drift, and vertically transmitted symbionts also compete for host colonization [51–54]. Thus, we should expect drift to generate heterogeneity in vertically transmitted infections [55,56] in the same manner we observed in our horizontal transmission experiments.

Based on our findings, we argue that the role of ecological drift is inadequately considered in host-microbe associations ranging from mutualistic to commensal to pathogenic [15,16,32,57,58]. Notably, as long as ecologically overlapping microbes are capable of colonizing a within-host niche, host benefit, partner choice and coevolutionary history play unexpectedly small roles in explaining the strain-level compositional variation in microbiomes observed between hosts [4,7,17,42,56,59–62], despite efforts to detect these effects [63–65]. On the other hand, niche differentiation and facilitation between taxa might be expected to promote stability (e.g., [66]). Whereas we studied ecological drift in a binary host-microbe symbiosis, in which interactions between strains are likely dominated by competition [7,38,51,67,68], multispecies communities may contain facilitative interactions such as signaling crosstalk, cross-feeding, metabolic division of labor, or complementary host-provisioning [10,69–76]. We postulate that facilitative interactions critical for host or microbial fitness resist stochastic loss of key, functionally non-redundant taxa, resulting in convergence in community composition. Nonetheless, previous work suggests that complex microbiomes associated with animals still assemble in a manner that conforms broadly to neutral expectations [23]. If transmission is so stochastic that it frequently decomposes multispecies communities, hosts that rely strongly on every unique function provided by each community member may evolve transmission mechanisms that minimize the effect of ecological drift (e.g., by increasing transmission bottleneck size [77]).

The pervasive effect of ecological drift suggests it may play a key but undervalued role in the evolution of specialized host-microbe symbiosis. First, ecological drift can override selection and maintain strain variation within a host population, by providing refugia for suboptimal or less competitive symbionts. In addition, by driving compositional variation between host-associated microbiomes, ecological drift can expose taxonomically or functionally distinctive strains and communities to selection [53,78–82]. If a distinctive microbiome can maintain its association with a particular host lineage, coevolution with the host may eventually occur. By simultaneously maintaining genetic variation among symbionts and generating heterogeneity in symbiont community composition, we argue that ecological drift could provide another explanation for the paradox of variation in host-microbe mutualisms [11].

Beyond its role in generating between-host heterogeneity, ecological drift also generates heterogeneity in symbiont populations within a host. In the squash bug, we found that gut crypts, a unique anatomical feature of the M4 symbiotic organ, generate heterogeneity by segregating strains into separate, discrete compartments within the same host. This has parallels in other symbiotic organs, including the crypts in the light organs and accessory nidamental glands of sepiolid squid [83–85], coralloid roots and root nodules in cycads and legumes [57,86,87], and pores in human skin [88]. Because we show that such compartmentalization acts even on isogenic cells, we propose that within-host population spatial structure, as with between-host population structure, is not adequately explained by either host selection or microbial competition, and is instead characterized by stochastic colonization of different crypts [57]. While spatial heterogeneity frequently emerges as a result of between-strain interactions within *in vitro* communities [89–93], here, the anatomy of a host forcefully imposes it even in the apparent absence of such interactions. How squash bugs and other multicellular hosts benefit from subjecting their symbiont populations to such elaborate compartmentalization remains an open question [94,95].

While our model system involves an extracellular symbiont, plasmids, organelles, and pathogens are also known to experience fine-scale population structure, with demonstrable effects on within-host evolution in addition to host fitness [81,96–102]. Among microbes, fine-scale population heterogeneity can drive the evolution of social cooperation within the host, by minimizing conflict with competitors (Kim et al., 2008; McNally et al., 2017), excluding social cheaters (Steinbach et al., 2020; van Gestel et al., 2014), and privatizing public goods (Kümmerli et al., 2009; McNally et al., 2017; van Gestel et al., 2014). To evaluate how symbiont evolution is affected by population heterogeneity within the host, more research remains to be done to characterize the spatial characteristics of within-host symbiont populations in a variety of animal and plant systems (e.g. [103,104]), as well as genetically manipulate between-strain interactions to understand the interplay between host-driven and microbe-driven determinants of structure [105].

Although we have discussed how ecological drift results in segregation of genetic variation within the host, we must address the most puzzling aspect of this finding in the context of our system, which is that the degree of within-host heterogeneity between single crypts depends on their location along the gut anterior-posterior axis. While posterior crypts are colonized by single isogenic strains, anterior crypts are always coinfected. Contrary to our expectations from *in vitro* systems [106], this pattern was sustained for all co-colonized individuals we observed, across four orders of magnitude in inoculum density. This suggests that *in vivo* colonization imposes unknown conditions distinct from *in vitro* colonization which creates structure within growing populations. We know almost nothing about symbiont colonization at the single-cell level in the squash bug that would result in this pattern, but we speculate that the host permits colonization by only a limited number of symbiont cells, resulting in these convergent results. While we consider it likely that this gradient in heterogeneity is a byproduct of symbiont dispersal within the M4 during early infection, we do not exclude the possibility that mixing multiple strains in the same compartment can provide unforeseen host benefits. Further study is necessary to ascertain whether coinfection within single crypts affects within-host symbiont evolution and host fitness, as predicted by others [107], or creates opportunities for horizontal gene transfer [108].

In this work, we illustrate the role of ecological drift in shaping symbiont host populations at multiple scales. Our findings highlight the effect of ecological drift during colonization by maintaining heterogeneity in symbiont populations both within and between hosts. We posit that ecological drift can overcome selection resulting from both host-microbe and microbe-microbe interactions, while also setting the stage for the evolution of these same processes. These results contribute to our understanding of the role that stochastic dynamics play in the assembly of ecological communities, even in ancient, highly specific host-microbe associations subjected to extensive host control [23].

## Acknowledgements

We thank the Gerardo, de Roode, Vega, and Levin lab members for helpful comments on this manuscript. In particular, we wish to thank Joselyne Chavez for supplying ASV diversity data from squash bugs from multiple field sites, Anthony Junker for important advice on microscopy and image analysis, Sandra Mendiola and Erik Edwards for maintaining squash bug lab colony stocks and squash plants, and Kayla Stoy, Justine Garcia, and Patrick Stillson for the isolation and genomic characterization of *Caballeronia* strains used in this study. We also thank Travis Wiles and Elena Wall for contributing the plasmids that made this project possible. This work was funded by Emory University and USDA NIFA 2019-67013-29371.

## Methods

### Study system

Squash bugs (*Anasa tristis*) were maintained on yellow crookneck squash plants (*Cucurbita pepo* ‘Goldstar’) in 1 ft^3^ mesh cages. Hatchlings were maintained on pieces of surface-sterilized organic zucchini in plastic rearing boxes, where they remain aposymbiotic (i.e., *Caballeronia*-free, though not necessarily free of other microbes). Hatchlings molt to the second instar, the life stage competent for symbiont colonization, after two days of feeding. Nymphs utilized in this experiment were typically one week old or less.

*Caballeronia* symbionts *C. sp.* SQ4a and *C. zhejiangensis* GA-OX1 were originally isolated from wild squash bugs at different localities in northeastern Georgia, USA. SQ4a and GA-OX1 form phenotypically very distinct colonies on nutrient agar (NA; 3 g/L yeast extract, 5 g/L peptone, 15 g/L agar) and are not closely related within the genus *Caballeronia* [35] (Fig S3). Cultures were typically grown on NA plates or in Luria Bertani (LB) Lennox broth with low salt (Sigma-Aldrich L3022), at 25°C. Unless otherwise stated, 2 mL broth cultures were initiated from colony picks of three to four day old colonies grown on NA at 25°C, and grown overnight with shaking at 200 rpm at 25°C.

### Whole genome-based classification of experimental symbiont strains

A whole genome-based phylogeny was constructed using RealPhy [109], with *Burkholderia cepacia* as the reference genome, using default settings except for a gap threshold of 0.1 and setting the model of evolution to GTR. Support values are bootstrap values based on 100 replicates. In addition to SQ4a and GA-OX1, *Caballeronia* strains A33M4c and IN-ML1 were also previously isolated from *A. tristis* [9,24]. SMT4a is a *Paraburkholderia terricola* soil isolate that can colonize *A. tristis* [24,35]. GenBank assemblies are as follows: GCF_023631065.1 (GA-OX1), GCF_022879815.1 (A33M4c), GCF_022627895.1 (*C. zhejiangensis*), GCF_023631085.1 (INML1), GCF_001544875.2 (*C. hypogeia*), GCF_000402035.1 (*C. insecticola*), GCF_023170545.1 (SQ4a), GCF_001544615.1 (*C. concitans*), GCF_902833485.1 (*C. glathei*), GCF_902859805.1 (*P. sediminicola*), GCF_022879555.1 (SMT4a) and GCA_009586235.1 (*B. cepacia*).

### Strain construction

The mini-Tn7 system [110] facilitates the stable, orientation-specific introduction of foreign DNA into bacterial genomes at a neutral intergenic site, *attTn7*, with minimal effects on phenotype and fitness *in vitro* [111–113]. Previously, our group introduced GFPmut3 into SQ4a using an archaic mini-Tn7 helper plasmid [36,114,115], prone to non-site specific genomic integration [116] and even whole plasmid integration. Moreover, poor fluorophore expression by the constitutive P_lac_-derived promoter P_A1/04/03_ in *Caballeronia* (also observed in *Burkholderia* [117]) necessitated the use of a higher-expressing mini-Tn7 construct for distinguishing fluorescent strains. Finally, Kikuchi & Fukatsu selected a spontaneously rifampicin resistant mutant as the basis for their labelling efforts [36]. To make readily distinguishable but otherwise isogenic symbiont strains without laboratory domestication, we genomically integrated a green fluorescent protein (sfGFP; henceforth GFP) and a red fluorescent protein (dTomato; henceforth RFP) into SQ4a and GA-OX1 using improved versions of previously developed mini-Tn7 vectors (Table 1) [110]. The conjugative *Escherichia coli* K12 strain SM10(λpir) harboring pTn7xKS-sfGFP or pTn7xKS-dTomato (Table 1), which were a generous gift from Travis Wiles [37], as well as an *E. coli* parent of the same strain harboring helper plasmid pTNS2 [110], were plated with SQ4a and GA-OX1 at high density on LB plates with salt (10 g/L tryptone, 5 g/L yeast extract, 10 g/L NaCl, 15 g/L agar). After 24-48 hours of incubation at 30°C, matings were harvested into LB Lennox low-salt broth with a lytic coliphage, T7, to eradicate *E. coli*. After further incubation for four hours at 30°C shaking at 200-225 rpm, cultures were plated on NA amended with 1 mM isopropyl-β-D-1-thiogalactoside (IPTG) and 10 µg/mL gentamicin to select for successful integrants. Colonies on selective plates were screened for fluorescence and frozen at -80°C as 20% v/v glycerol stocks.

**Table 1.** Strains and plasmids used in this study.

| Strains | Description | Source/Reference |
| --- | --- | --- |
| *Escherichia coli* SM10(λpir) | recA::RP4-2-TcR::Mu, pir+ conjugative strain | Travis Wiles [118,119] |
| *Caballeronia zhejiangensis* GA-OX1 | Wild *Anasa tristis* isolate, Oxford College Organic Farm | [9] |
| *Caballeronia* sp. nr. *concitans* SQ4a | Isolated from a wild-caught *Anasa tristis* held in captivity; originally collected at Oakhurst Community Garden | [24] |
| Plasmids | Description | Source/Reference |
| pTNS2 | R6K $\gamma$ oriV, AmpR, tnsABCD | Travis Wiles [110] |
| pTn7xKS-sfGFP | colE1 oriV, mini-Tn7T-GmR-sfGFP, AmpR, lacI P<sub>tac</sub> hokB ghoT tisB | Travis Wiles [37] |
| pTn7xKS-dTomato | colE1 oriV, mini-Tn7T-GmR-dTomato, AmpR, lacI P<sub>tac</sub> hokB ghoT tisB | Travis Wiles [37] |

To confirm stability of fluorophore expression, newly constructed strains were streaked on NA plates and visually assessed for fluorescence after two days. To confirm site- and orientation-specificity of mini-Tn7-GmR integration, we ran PCR to amplify the fragment between the endogenous *glmS* gene (GA-OX1: 5’ AGGCGCGTTGAAGCTCAAGG 3’; SQ4a: 5’ CGCTGGAGCCGCAAATCATC 3’) and the inserted *aacC* gentamicin resistance marker (aacC-83F: 5’ GTATGCGCTCACGCAACTGG 3’). We did not screen for insertion at additional sites, as to our knowledge the strains of *Caballeronia* we used have only one *attTn7* site.

### Competition assays *in vitro*

During the summer growing season, squash bugs feed on macerated cell contents, xylem, and phloem in tissues from squash plants and fruits [120,121]. To replicate microbial competition in this environment, competition assays in liquid culture were conducted in filter-sterilized zucchini squash extract. In short, juice from organic zucchini fruits was extracted in a juicer, combined, and filtered to remove large suspended particles. This filtrate was then centrifuged at 10,000 xg for 3 hours to pellet suspended particles, then filter-sterilized through a 0.2 µm filter and stored at -20°C.

To initiate competition assays, GA-OX1 and SQ4a, labelled with sfGFP or dTomato as described above, were initially streaked from frozen 20% glycerol stocks onto NA plates and incubated at 30°C for 48 hours. Individual colonies from each plate were inoculated into 3 mL of LB media and incubated in a shaking incubator (New Brunswick Scientific Excella E25) at 30°C for 12 hours with shaking at 225 rpm. All cultures were equalized to an optical density (OD) of 1.0 by adding 100 µL of each culture to a 96 well plate and taking readings with a Synergy HTX multimode plate reader. The equalized cultures were spun down with an Eppendorf centrifuge 5424 R and washed with one mL of 1X phosphate buffered saline (PBS) three times.

Monocultures of GA-OX1 and SQ4a were combined to form counterlabelled self vs. self and self vs. competitor co-cultures, for a total of four combinations. Self vs. self co-cultures contained the same *Caballeronia* strain, differing only in the fluorescent protein, while self vs. competitor co-cultures contained different *Caballeronia* strains, also differing in the fluorescent protein. All co-cultures were subsequently incubated for 24 hours at 30°C with shaking at 225 rpm. As described above, co-cultures were dilution plated on NA and incubated a further 20-24 hours, and single colonies containing each fluorophore were distinguished and counted under a dissecting scope. We confirmed that SQ4a and GA-OX1 do not appear to inhibit each other intensely on NA plates, suggesting that plating co-cultures on NA provides an unbiased count for both competitors (Fig S5).

### Competition assays *in vivo*

The generalized protocol for competition assays with varying transmission bottlenecks *in vivo* is presented in Fig 2A. GFP- and RFP-labelled *Caballeronia* strains were streaked out from glycerol stocks onto NA and incubated at 25°C for at least three days. To initiate liquid cultures, single colonies were picked into two mL LB in glass tubes and incubated at 25°C with shaking at 200 rpm; to account for different growth rates between SQ4a and GA-OX1, a glass inoculating loop was used to pick up an entire colony of SQ4a, while a p10 micropipettor tip was used to extract a small plug from part of a single GA-OX1 colony.

To prepare inocula for feeding, cultured bacterial cells were washed to remove LB. Two hundred µL of culture was spun down at 10,000 x g at 4°C for 2 min. The supernatant was removed, and the pellet was resuspended in 1000 µL 1X PBS. After a second centrifugation, the pellet was resuspended in 1X PBS. For quality assurance, tenfold serial dilutions were carried out using 30 µL washed cells in 270 µL PBS in a 96-well plate. Fifty µL each of the washed GFP- and RFP-labelled strains were then diluted into 400 µL of a complex feeding solution (a 1:1 mixture of filter-sterilized zucchini squash extract and PBS; for neutral competition trials) or a defined feeding solution (2% m/v glucose 10% v/v PBS; for interspecies competition trials), containing 20 µL of a nontoxic blue dye (1 mM erioglaucine disodium). We found that the defined feeding solution improved nymphal feeding response and better prevented bacterial population growth during the inoculation time window, with minimal impact on our experimental results (Fig S4). Nymphs previously starved overnight for 15-25 hours in clean plastic rearing boxes (7 cm X 7 cm X 3 cm) were supplied with 120 µL of a single inoculum treatment blotted on quartered sectors of 55 mm-diameter qualitative filter paper (Advantec MFS N015.5CM). Nymphs were then allowed to wander and feed *ad libitum* for 2-3 hours. After this brief inoculation period, nymphs were housed singly in 24 well plates with small pieces of organic zucchini to develop for three days. Just before and after the inoculation period, inocula were serially diluted as above to quantify the concentration of each strain and ensure that no substantial growth or death of either strain occurred during the inoculation period (Fig S6).

On the fourth day after inoculation, nymphs were killed in 70% denatured ethanol, surface sterilized in 10% bleach for 5-10 minutes, washed off again in 70% ethanol, and immersed in ∼20 µL droplets of 1xPBS. Whole nymphs, when applicable, were imaged on a Olympus SZX16 stereomicroscope with an Olympus XM10 monochrome camera and Olympus cellSens Standard software ver. 1.13. Nymphs were immersed in a shallow volume of PBS in 6 cm plastic petri dishes, and images were taken in darkfield (30 ms exposure 11.4 dB gain), brightfield (autoexposure, 11.4 dB gain), a GFP channel (autoexposure, 18 dB gain), and a RFP channel (autoexposure, 11.4 dB gain). Darkfield and brightfield images were merged in FIJI version 1.54f using the Image Calculator plugin, and the result was then merged with the GFP channel, RFP channel, or both. M4s were individually dissected from nymphs, and the degree of green and red colonization was qualitatively estimated under a fluorescent microscope. Each M4 was then held in 300 µL 1x PBS in Eppendorf tube and crushed with a sterile micropestle. Thirty µL of homogenate was serially diluted in 270 µL PBS and immediately dilution plated onto NA. Plates were then incubated at 30°C for about 24 hours. Counts of GFP and RFP fluorescent colonies were recorded after refrigeration at 4°C for at least 24 hours to enhance fluorescent protein expression. The count data of GFP and RFP colonies yielded by our sampling procedure almost always reflected our qualitative observations of GFP and RFP colonization inside the M4, suggesting that our data accurately represent the colonization state within live insects.

#### Microscopy of Within-Host Symbiont Populations

Bugs were inoculated and allowed to develop for four days as described above with ∼60 and 931,000 CFU/µL inocula containing GA-OX1 GFP and RFP. From each bug, the whole gut was dissected in a 20-30 µL droplet of PBS in a 30 mm diameter plastic dish. The M4 was stretched out to its full length, and straightened out as much as possible by severing tracheoles associated with the crypts and flipping the M4 over to minimize the number of twists in the M4. This was critical to minimize aberrations in fluorescence intensity and colocalization due to overlap between multiple crypts. The M4 was anchored at the posterior end by the tip of the bug abdomen and at the anterior end by the M1-M3 sections of the midgut, and cleaned several times by pipetting off debris, fat body, and hemocytes with clean PBS. Finally, the whole preparation was re-immersed in 2550 µL of PBS, to which 1 µL of M9 buffer containing 1% Triton-X100 was added to aid the spreading of the droplet.

Gut preparations, which degrade or dry rapidly, were imaged as soon as possible. Tilescan images were taken using a Leica DMi8 inverted widefield light microscope with a Leica DFC9000 GT fluorescence camera and Leica Application Suite X ver. 3.4.2.18368 software. Automated tilescans were taken with a 10X objective lens with brightfield, DIC, GFP, and RFP channels. Fluorescent channels were established by filter sets. The GFP channel was set to: bandpass filter 470/40 nm emission, dichroic mirror 495 nm, emission 525/50 nm. The dsRed channel was set to 546/11 nm excitation, dichroic mirror 560 nm, 630/75 nm emission. As each sample is unique, care was taken to set GFP and RFP channel exposure times manually according to the most intense pixels in the entire M4 (usually in the posterior crypts), to minimize signal saturation in any part of the preparation. Due to the convoluted shape of the M4, images were taken with and without autofocus, and stitched images were visually assessed to determine which images were more useful. The repetitive structure of the M4, composed of nearly identically sized, regularly spaced crypts, also necessitated a lower overlap value between tiles for tilescans, as low as 2%. LAS X software was used to merge tiles from tilescans without smoothing for quantitative analysis.

### Statistical analysis

All statistical analyses were conducted in R version 4.1.1, and the R package ggplot2 (version 3.4.2) was used for all data visualization. Because multiple trials were run for inoculation experiments, and some trials recovered very low numbers of infected nymphs, we binned nymphs from multiple trials into discrete treatment groups, based on inoculum size, for analysis. For neutral competition experiments, which utilized isogenic GA-OX1 GFP and RFP, we measured the proportion of GA-OX1 RFP extracted from each host. For interspecies competition experiments, utilizing different combinations of SQ4a and GA-OX1, we measured the proportion of GA-OX1, the superior competitor, compared to the sum of all green and red fluorescent colonies extracted from each host.

Raw colony counts of each fluorescent strain recovered from each individual insect were converted into proportions for analysis and visualization. To quantify between-host heterogeneity in symbiont colonization for each inoculation treatment, we calculated a bimodality coefficient [33] using the R package mousetrap (version 3.2.0), as well as the population variance, for each inoculation treatment. We also independently validated this approach using a species-level Fst measure as described previously for quantifying compositional variation among plant communities [21], which is based on the classical Fst measure for quantifying genetic differentiation between populations [122]. Using the R package diptest (version 0.76-0), we also implemented Hartigan’s dip test [123], which calculates a dip statistic (Tables S1-S4) based on the shape of the cumulative distribution function of a dataset. In addition to the dip statistic, we also calculated three p-values for each treatment to assess deviation from normality of the distribution of GA-OX1 compositional variation. We considered a distribution to deviate from unimodality if p-values from the dip test repeatedly fell below the threshold of 0.05.

To quantify within-host spatial structure, a linear region of interest (ROI) was sampled from one complete row of crypts from each sample to obtain GFP and RFP intensities. Crossover of the ROI from one side of the M4 to the next was occasionally necessary to follow that row through each twist of the M4. The identical ROI was translated to obtain GFP and RFP intensities from the empty background immediately adjacent to the crypts. GFP and RFP intensities at each point along the M4 were normalized by subtracting the background signal from the same point outside the M4. For the RFP channel, the difference between crypt and background signal was occasionally less than 0; in these rare cases the normalized RFP intensity at that point was assigned a value of 1. The log-transformed ratio of RFP to GFP intensity was measured from each pixel along the ROI. In addition, the variance in this value was calculated by iteratively sampling pixels from within a sliding interval 10% of the length of the ROI.

## Supporting information

**S1 Fig. Unaggregated infection outcomes resulting from neutral competition during isogenic co-colonization.**

A) Relative GA-OX1 RFP abundance within inocula used in isogenic colonization trials. Vertical lines demarcate which trials were aggregated for analysis in Fig 1. Black X marks indicate the inoculum density, and the percent relative abundance of GA-OX1 RFP, in each trial. The dotted horizontal line represents the average relative abundance of GA-OX1 RFP (46.5%) across all trials.

B) Variable colonization outcomes associated with different transmission bottleneck sizes in isogenic co-inoculation, disaggregated from Fig 2D. Gray X marks indicate the inoculum density, and the percent relative abundance of GA-OX1 RFP, in each trial. Points indicate successfully colonized nymphs associated with each inoculation trial, and the color of each point and its position along the y-axis represent the percent relative abundance of GA-OX1 RFP colonies among all fluorescent colonies recovered from each nymph. Magenta points represent insects containing only RFP colonies, green points represent insects containing only GFP colonies, and faded magenta/green colonies are co-colonized. Note that multiple points overlap

C) Bimodality coefficients calculated from unaggregated trials. Bimodality coefficients (black points) calculated from data in panel B. The 0.555 threshold (marked with a dotted line) indicates the bimodality coefficient expected from a uniform distribution.

**S2 Fig. Isogenic coinfections of *A. tristis* nymphs over five orders of magnitude in inoculum density.**

A) Fluorescence images of nymphs from a single cohort colonized with different densities of GA-OX1 sfGFP and GA-OX1 RFP, ranging from 10^1^ to 10^6^ CFU/µL. Nymphs inoculated with only GA-OX1 sfGFP or only GA-OX1 RFP serve as controls (top 2 rows). Note that the red fluorescent protein dTomato appears more fluorescent in whole-body preparations of nymphs than the green fluorescent protein sfGFP, due to increased absorbance of green light by living tissue [124] and the high stability of free dTomato under physiological conditions.

B) Infection outcomes of bugs associated with these trials, disaggregated from Fig 1D.

C) Bimodality coefficients calculated from unaggregated trials.

**S3 Fig. Symbiont strains SQ4a and GA-OX1 represent distinct clades within the genus *Caballeronia*.** Whole genome-based phylogeny of selected species and *Anasa tristis* symbionts representing major clades within the genus *Caballeronia*, as defined by Peeters *et al.* (2016) [26], including the experimental strains *C. sp. nr. concitans* SQ4a and *C. zhejiangensis* GA-OX1.

**S4 Fig. Different combinations of competing GA-OX1 and SQ4a yield qualitatively similar responses to increasing ecological drift in transmission.**

A) Variable colonization outcomes associated with different transmission bottleneck sizes in two-species co-inoculation, using *C. sp. nr. concitans* SQ4a sfGFP and *C. zhejiangensis* GA-OX1 RFP. Gray X marks indicate the mean percent GA-OX1 RFP associated with each inoculum treatment, ranging from approximately 10^0^ to 10^5^ CFU/µL. Points represent individual nymphs, and the color of each point and its position along the y-axis represent the percent relative abundance of GA-OX1 RFP colonies among all fluorescent colonies recovered from each nymph. Magenta points represent nymphs from which only GA-OX1 RFP colonies were recovered, green points represent nymphs from which only SQ4a sfGFP colonies were recovered, and faded magenta/green points represent coinfected nymphs. Violin plots associated with each treatment depict the shape of the distribution in relative GA-OX1 RFP abundance. Below each violin plot, the success rate of colonization is indicated, as the number of nymphs that were successfully colonized with *Caballeronia* out of all nymphs sampled. These values were not recorded for the 10^0^-10^1^ treatment and thus omitted. Asterisks indicate significantly multimodal infection outcomes as determined by Hartigan’s dip test, at a significance level of p < 0.05.

B) Bimodality coefficients calculated from results in panel A. Colonization is bimodal (bimodality coefficient > 0.555) across several orders of magnitude of inoculum density. The 0.555 threshold (marked with a dotted line) indicates the bimodality coefficient associated with a uniform distribution.

C) Variable colonization outcomes associated with different transmission bottleneck sizes in two-species co-inoculation, using *C. sp. nr. concitans* SQ4a RFP and *C. zhejiangensis* GA-OX1 sfGFP. Gray X marks indicate the mean percent GA-OX1 GFP associated with each inoculum treatment, ranging from 10^1^ to 10^5^ CFU/µL. Points represent individual nymphs, and the color of each point and its position along the y-axis represent the percent relative abundance of GA-OX1 GFP colonies among all fluorescent colonies recovered from each nymph. Magenta points represent nymphs from which only SQ4a RFP colonies were recovered, green points represent nymphs from which only GA-OX1 sfGFP colonies were recovered, and faded magenta/green points represent coinfected nymphs. Violin plots associated with each treatment depict the shape of the distribution in relative GA-OX1 GFP abundance. Below each violin plot, the success rate of colonization is indicated, as the number of nymphs that were successfully colonized with *Caballeronia* out of all nymphs sampled. Asterisks indicate significantly multimodal infection outcomes as determined by Hartigan’s dip test, at a significance level of p < 0.05.

D) Bimodality coefficients calculated from results in panel C. Colonization is bimodal (bimodality coefficient > 0.555) across several orders of magnitude of inoculum density. The 0.555 threshold (marked with a dotted line) indicates the bimodality coefficient associated with a uniform distribution.

**S5 Fig. GA-OX1 and SQ4a do not exhibit strong inhibition on nutrient agar.**

Spots of GA-OX1 RFP and SQ4a sfGFP plated side-by-side at high and low densities on nutrient agar.

A) A dense culture of GA-OX1 RFP spotted adjacent to single SQ4a sfGFP colonies.

B) A dense culture of SQ4a sfGFP spotted adjacent to single SQ4a sfGFP colonies.

C) A dense culture of SQ4a sfGFP spotted adjacent to single GA-OX1 RFP colonies.

**S6 Fig. Changes in titer of each strain during all inoculation trials.**

Bacterial strain titers before and after inoculation trials.

A) GA-OX1 GFP and RFP (cf Figs 2B, 2C, and S1)

B) SQ4a GFPmut3 and GA-OX1 RFP (cf Figs 3C and 3D)

C) SQ4a sfGFP and GA-OX1 RFP (cf Figs S4A and S4B)

D) GA-OX1 sfGFP and SQ4a RFP(cf Figs S4C and S4D)

